# Structural Analysis of Spike Protein Mutations in the SARS-CoV-2 P.3 Variant

**DOI:** 10.1101/2021.03.06.434059

**Authors:** Neil Andrew D. Bascos, Denise Mirano-Bascos, Cynthia P. Saloma

## Abstract

A SARS-CoV-2 lineage designated as P.3 with multiple signature mutations in the Spike protein region was recently reported with cases from the Central Visayas Region of the Philippines. Whole genome sequencing revealed that the 33 samples under this lineage all contain the E484K, N501Y, and P681H Spike mutations previously found in variants of concern (VOC) such as the B.1.351, the P.1 and B.1.1.7 variants first reported in South Africa, Brazil, and the United Kingdom, respectively. The possible implications of the mutations found in the Spike protein of P.3 were analyzed for their potential effects on structure, stability, and molecular surface character. The analysis suggests that these mutations could significantly impact the possible interactions of the Spike protein with the ACE2 receptor and neutralizing antibodies, and warrants further clinical investigation. Some of the mutations affecting the N and C terminal domains may have effects on Spike monomer and trimer stability. This report provides insights on relevant targets for the design of future diagnostics, therapeutics and vaccines against the evolving SARS-CoV-2 variants in the Philippines.

## INTRODUCTION

SARS-CoV-2 is a single-stranded RNA virus belonging to the coronavirus family. The virus was first identified in January 2020 (WHO, 2020; Wang et al., 2020) and is the causative agent of the ongoing COVID-19 pandemic worldwide. Continued transmission from person-to-person and prolonged disease in some individuals have resulted in the emergence and detection of several variants of the virus towards the last quarter of 2020. These include the highly publicized UK variant, B.1.1.7 (Tang, Tambyah and Hui, 2020) which has increased transmissibility, and the South African variant B.1.351 (Tegally et al., 2020) which has been shown to be able to evade neutralization by some antibodies raised against the original SARS-CoV-2 (Weisblum et al., 2020; Wibmer et al., 2021, Xie et al., 2021). Several signature mutations have been reported for lineage P.3 of SARS-CoV-2 viruses from the Philippines (EpiCoVTM database of the Global Initiative for Sharing All Influenza Data (GISAID) with accession numbers EPI_ISL_1122426 to EPI_ISL_ 1122458).

The presence of 13 lineage-defining mutations with seven (7) occurring in the Spike protein region hint at the development of a unique SARS-CoV-2 variant within the country. Here, we analyze the potential significance of the reported mutations within the infection-relevant Spike protein in terms of its effects on structure and potential interactions. Analyses of the structural properties and their implications were done using *in silico* methods focused on detecting changes in intramolecular, and intermolecular interactions, as well as effects on protein surface character.

## MATERIALS AND METHODS

### SARS-CoV2 Sequences

The SARS-CoV-2 sequences were sourced from the GISAID database with EpiCov accession codes EPI_ISL_1122426 to EPI_ISL_ 1122458 and based on their reported lineage-defining mutations (Tablizo et al., 2021). A summary of all the mutations in the Spike protein present in the variants is presented in Table 1.

**Table 1.**
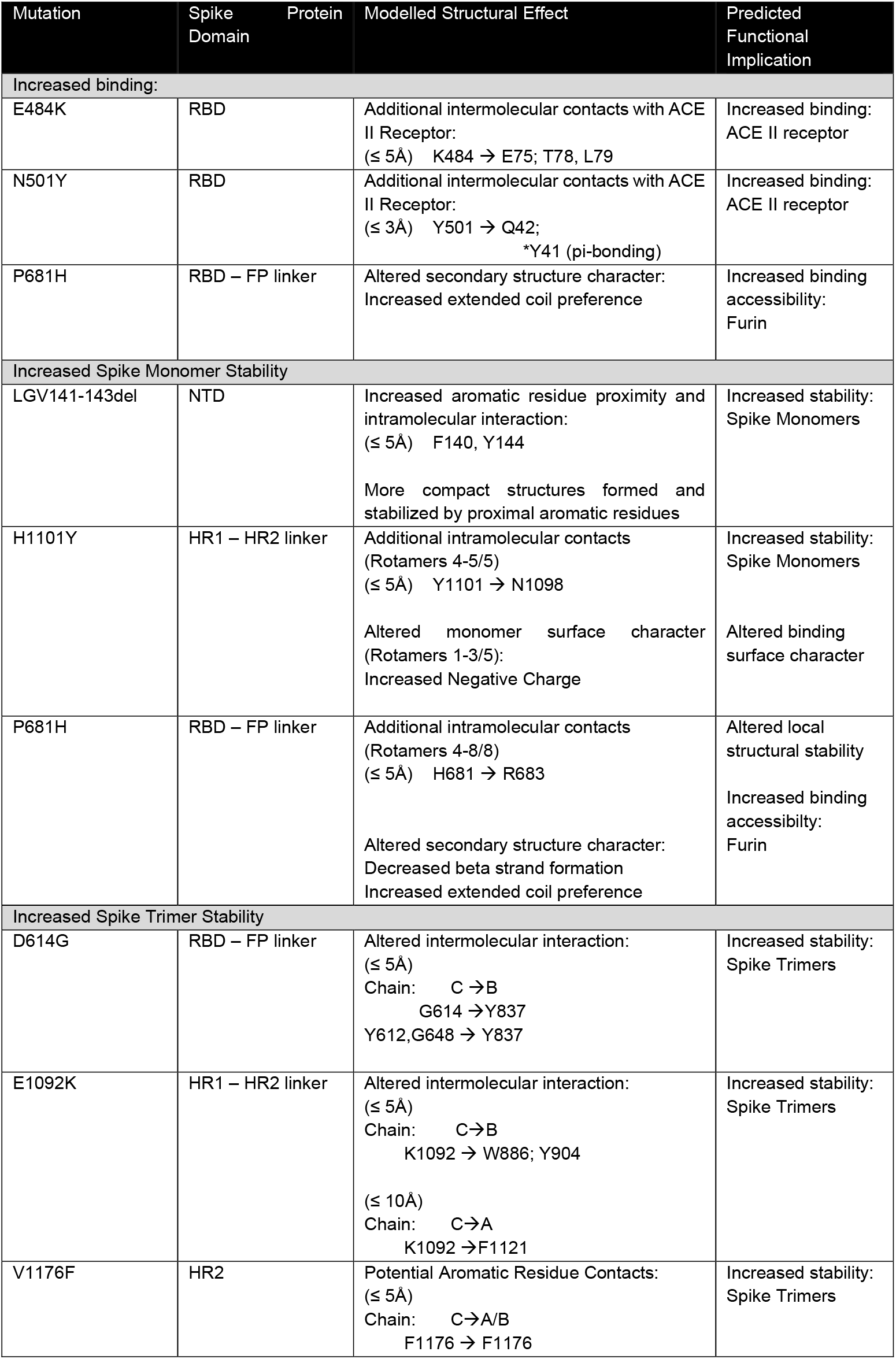
Summary of Signature Spike Protein Mutations in the Emergent SARS-CoV-2 Variant

### Molecular Modelling

Molecular structures for the SARS-CoV-2 Spike proteins were acquired from the iTASSER server of the Zhang Lab (2020). Models of different SARS-CoV-2 virus proteins (e.g. Spike protein, RdRp, ORF-8, etc.) were downloaded in monomeric, trimeric (S.pdb), and receptor bound forms (e.g. S_ACE2.pdb) and analyzed using several molecular modelling software. iTASSER generated models of documented SARS-CoV-2 variants of concern (B.1.1.7; B.1.351 and P.1) were also used in the analysis.

Basic structural analysis and residue contact identification was done using the DeepView modeller (Guex and Peitsch, 1997). Models of the different domains of the SARS-CoV-2 Spike protein were generated based on the domain specifications listed by Huang et al. (2020).

### Mutant Protein Model Generation

Several modelling software were used to generate models with the mutated residues. The mutate function of DeepView was used to generate the mutant protein models for variants with amino acid substitutions (e.g. N501Y, E484K, etc.). When several rotamer forms are possible for the generated mutant residue, all positions were observed for potential contact alterations.

In some cases, the mutations did not result in altered contact but in differences in the charge/character of the molecule’s surface. Rendering of molecular surfaces was done using VMD 1.9.3 (Humphrey et al., 1996; https://www.ks.uiuc.edu/) to illustrate nonpolar, and charged (positive / negative) regions on the molecular model. Models for mutations that involved residue deletions (e.g. LGV141-143del, Y144del) were made using RaptorX (Källberg et al., 2012; raptorx.uchicago.edu).

## RESULTS

The SARS-CoV-2 virus continues to accumulate mutations as it is transmitted from host to host as part of its normal process of evolution. Thousands of mutations in the virus have already been detected with the unprecedented pace of virus genomic sequencing being done worldwide to study and track its spread. This is exemplified by the growth of data sharing seen in the GISAID EpiCoV database which has about 699,000 deposited sequences of SARS-CoV-2 as of March 6, 2021. Many of the mutations in the coronavirus are inconsequential (e.g., synonymous mutations), but when multiple mutations in the Spike region are detected in the context of an increasing epidemiological curve, this elicits interest due to its potential clinical and public health implications. Here, we modelled the possible structural consequences of the multiple Spike protein mutations in the P.3 lineage detected in the Central Visayas region of the Philippines. The mutations were classified based on their predicted effects on Spike protein interactions both to the ACE II receptor and other co-interacting proteins, such as neutralizing antibodies; and effects on monomer and trimer stability. These predicted effects are summarized in Table 1.

### Presence of the E484K and N501Y mutations strengthen Spike protein – ACE II receptor interaction

Perhaps the most profound changes in Spike protein function would be those that affect its interaction with the ACE II receptor. Successful interaction with the receptor ultimately leads to the facilitation of viral entry and infection. Changes that lead to increased interaction between the Spike protein’s RBD and the ACE II receptor would be associated with increased probability of viral infection.

Both of the residue substitutions found in the RBD for the new variant (i.e. E484 and N501) are located at the interface with the ACE II receptor. Both of the observed mutations for these residues (i.e. E484K and N501Y) are predicted to increase interactions with the ACE II receptor (Figure 1). These mutations have been previously studied both *in silico* and in vitro for effects on infectivity (Honying, et al., 2020; Starr et al., 2020; Brouwer, et al. 2020; Zhang, et al., 2020.

**Figure 1.**
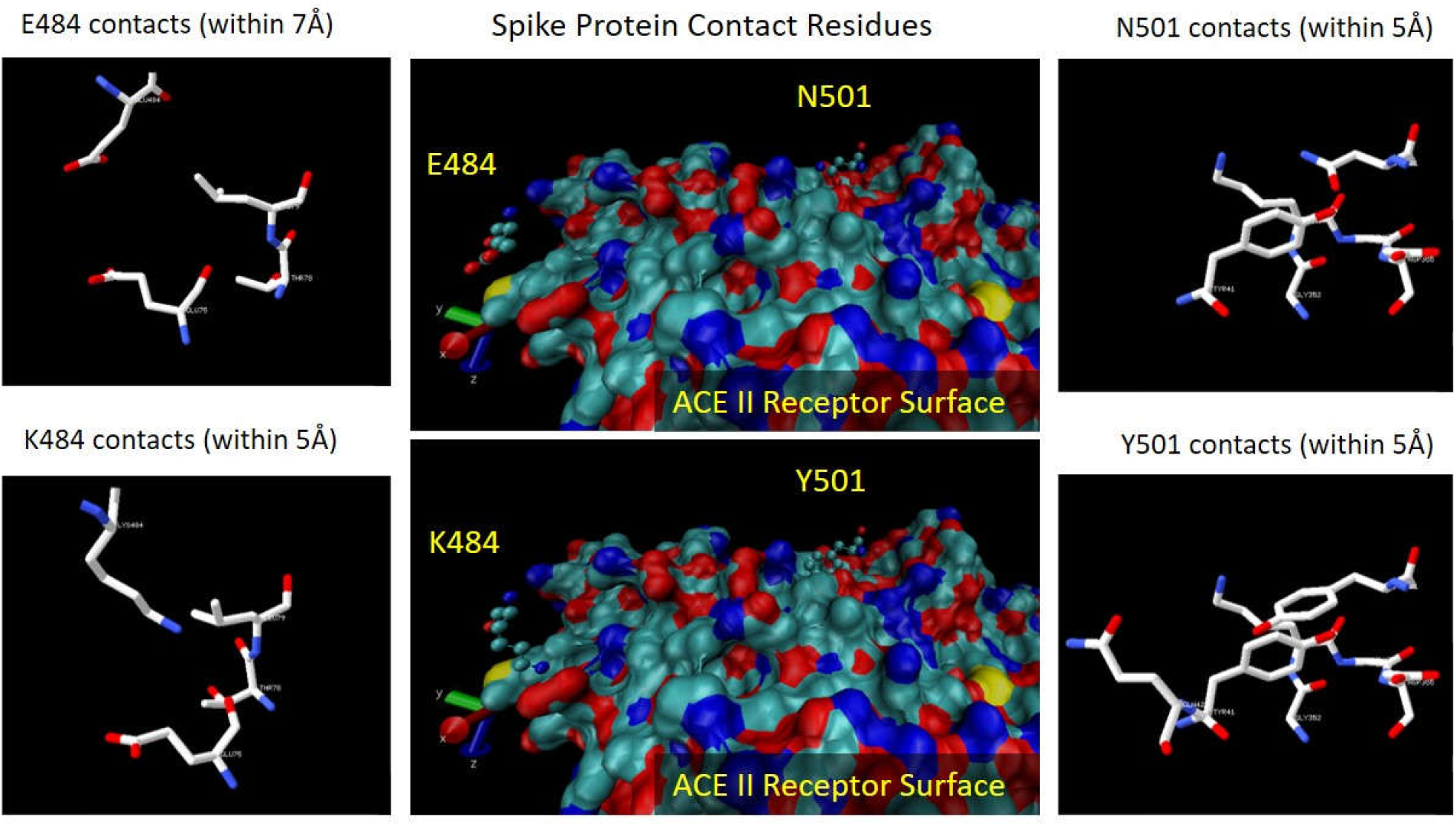
Contacts with ACE II Receptor Surface The surface of the ACE II receptor that interacts with the Spike Protein RBD is presented in the central boxes, with residues colored based on their charge characteristics (Red, negative; Blue, positive; and Cyan, non-polar). Yellow surfaces denote the presence of Sulfur atoms. Both E484 and N501, depicted as ball and stick models, are seen to be placed close to the ACE II surface, with N501 fitting within a complementary charged pocket. E484’s negatively-charged side-chain is shown to be oriented away from the nonpolar surface. Replacement of these residues with K and Y, respectively, result in additional contacts with the ACE II receptor surface. K484 gains better access and complementarity with the negative charged surface residues. Y501 is able to contact more residues, and may potentially interact with Y41 on ACE II through pi-bonds.

Glutamate 484 (E484) is located at the edge of the RBD-ACE II interface. Interestingly, the charge character of E484 is not complementary to its proximal ACE II receptor surface, and it was not observed to contact ACE II receptor residues within a 5Å cut-off distance. The first contacts were observed to occur at 6Å from E484. This was changed with the conversion into Lysine (E484K). The elongated sidechain of K484 resulted in new contacts with compatible residues in the ACE II receptor (E75, T78, L79) within 5Å. The E484K substitution is predicted to result in additional contacts between the Spike protein RBD and the ACE II receptor similar to other reported variants of concern flagged by the WHO which both contain these amino acid alterations in the Spike protein region (Ramanathan, et al., 2021). Residue contacts between the RBD and ACE II were previously documented in a crystallized structure of the two (2) structural domains (Wang et al. 2020, PDBID 6LZG). In this form, E484 was observed to interact with K31 of the ACE II receptor via a salt bridge. This salt bridge was predicted to be lost with the molecular dynamics simulation studies conducted by Cheng et al. on the South African 501.V2 variant (Cheng et al., 2021). It must be noted that this salt bridge is also not present in the iTASSER modelled structure for the Spike protein trimer upon which our models were based, suggesting the possibility of different conformations attained within crystallizing conditions, and the buffered environments which the modelling simulations approximate. In our models, the E484K mutation would allow contact with an adjacent glutamate residue on the ACE II receptor (E75), providing additional stability to the RBD-ACE II interaction.

From a structural standpoint, increased efficiency of binding of the E484K mutant to the ACE2 inhibitor diminishes the chance of antibodies accessing potentially neutralizing sites. This shift of the system’s equilibrium towards more Spike-ACE2 binding implies a decrease in effective Spike-antibody binding. The E484K mutation has been found to diminish antibody neutralization potentials in experimental studies (Weisblum et al., 2020; Wibmer et al., 2021, Xie et al., 2021). The efficacy of the Novavax vaccine has also been observed to be significantly reduced in cases caused by viruses with this particular mutation (Mahase, 2021) Thus, like other SARS-CoV-2 mutations of interest, the presence of these mutations in the emergent Philippine variant (P.3) is of particular concern, and warrants close surveillance.

Asparagine 501 (N501) is found in the center of the RBD-ACE II interface. The charged character of N501 is observed to complement the binding surface on ACE II. Asparagine 501’s location allows it to interact with several residues on the receptor surface (Y41, G352, K353, G354, D355) at a cut-off distance of 5Å. Of these, Y41 is closest, interacting within 3Å. The conversion of residue 501 to Tyrosine (Y501) allows additional contacts to be made. An interaction with Q42 is added within the 5Å cutoff, and K353 is added within 3Å. Interestingly, the conversion into Y501 also allows the formation of a pi bond interaction with Y41 (Figure 1). The interaction of Y501 and Y41 aromatic rings would be expected to stabilize the RBD-ACE II receptor interaction.

The N501Y mutation has been documented in variants from the UK, Brazil, and South Africa. This particular mutation has been hypothesized to increase the transmission of SARS-CoV-2 by anywhere between 10 - 75% (Zhao, et al., 2021, Leung et al, 2021). The increase in cases in areas where the N501Y mutation has been identified supports this hypothesis necessitating measures to control its transmission and to implement enhanced contact tracing protocols up to the third generation.

The structures of several SARS-CoV-2 variants have been modelled and are available through iTASSER (Zhang lab, 2020). The three (3) variants of concern (VOCs): B1.1.7, B.1.351 and P.1, all harbor the N501Y mutation whereas the latter two (2) VOCs possess both the E484K and N501Y substitutions. Both of these RBD mutations are observed in P.3 viruses.

To evaluate the effects of these mutations on RBD: ACE II binding, accessible residues in the ACE II receptor for either Y501, or K484 were identified. It must be noted that the available models in iTASSER for the SARS-CoV-2 variants only include a few helices to represent the ACE II receptor (Figure 2A). However, by fitting the RBDs of the variant structures unto the complete Spike-protein:ACE II receptor structure (S_ACE2.pdb; Zhang lab, 2020), relevant residue contacts may be observed. Identification of accessible residues in the ACE II receptor for the RBD: K484, and N501 coincide with the predicted contacts of the P.3 variant models (Figures 1 and 2A). Potential rotamer conformations of K484 place it within 5.37Å of E75 in the ACE II receptor. N501 is also observed to contact the ACE II receptor through Y41 within 3.06 Å. The observation of these additional contacts in the modelled variant protein (Figure 2A) provides supporting evidence for the compounded effects of these two (2) mutations for stabilizing RBD: ACE II interactions, and ultimately enhancing infective potential of these emergent variants of SARS-CoV-2.

**Figure 2.**
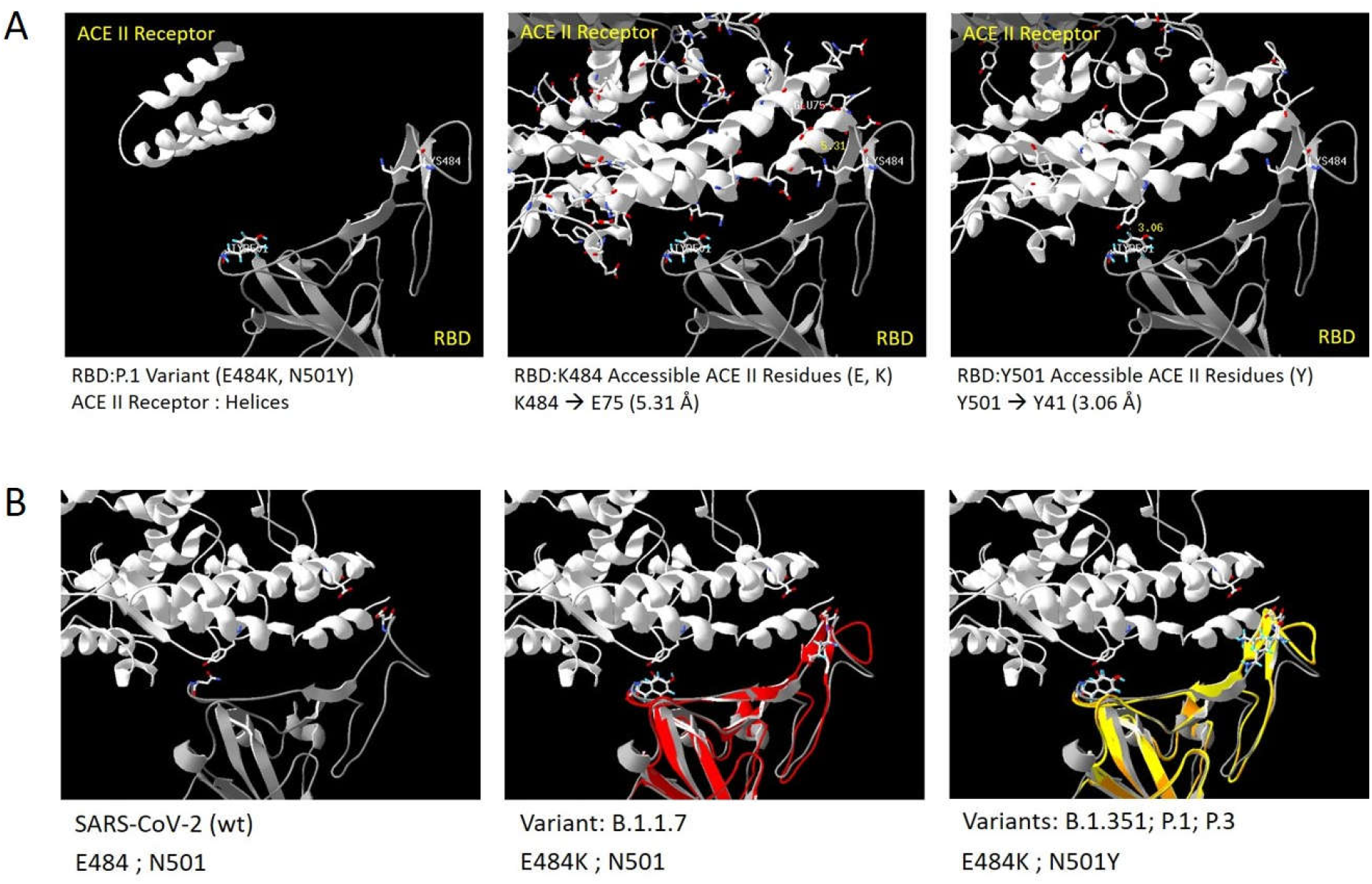
SARS-CoV-2 variant Contacts with ACE II Receptor (A) A model structure for the P.1 Variant of SARS-CoV-2 contains both E484K and N501Y mutations. (Left) The model in iTASSER (P.1 model, open.pdb; Zhang lab, 2020) does not contain the interfacial contacts between the Spike protein RBD (gray) and the ACE II receptor (white). Fitting the ACE II receptor domain (white) from the S_ACE2.pdb structure (Zhang lab, 2020) into the model allows the observation of the interacting residues in both domains. (Center) The glutamate and lysine residues, which may form salt bridges, are shown as linear models. E75 in the ACE II receptor is shown to be accessible to K484. One rotamer conformation is observed to come within 5.31Å of E75. (Right) All ACE II Tyrosine residues were rendered in linear models to show their location. Tyrosine 41 is observed to be accessible to Y501 (separation distance: 3.06Å) in this model for the P.1. variant. (B) The interface of the Spike protein RBD and the ACE II receptor has been the site of amino acid substitutions in the different SARS-CoV-2 variants. The emergent variant from the UK (B.1.1.7) was observed to have an N501Y mutation. This mutation has been observed in succeeding identified variants from South Africa (B.1.351), Brazil (P.1) and now, the Philippines (Ph-B.1.1.28). In addition, a E484K substitution is observed in the 3 latter variants. The relative positions of the wt and variant RBD models are overlaid in the figure (WT, gray; UK, red; South Africa, orange; Brazil, yellow). Both N501Y and E484K are predicted to provide additional contacts between the RBD and ACE II receptor. N501 is observed to exist in a stable conformation across the 3 variants shown. K484 is observed to have multiple rotamer conformations, some of which may access E75 in the ACE II receptor.

The appearance of the N501Y and E484K mutations in several variants all over the world (Figure 2B), presumably from independent events, highlights the selective pressure that is upon these particular residues of the Spike protein and points to a possible convergent evolution of the virus at these sites. It will be prudent to consider that the design of future vaccines against SARS-CoV-2 should take both these mutations into account.

### Additional mutations that could impact Spike protein surface character

In addition to altered interactions with the ACE II receptor, changes in the other binding surfaces of the Spike protein may also prove important. Changes of this sort have shown clinical relevance, as seen with the Y144del mutation associated with the B.1.1.7 UK variant. This mutation was seen to result in a decreased efficiency of neutralization by antibodies (McCallum et al. 2021). Analysis of the Spike protein surface at the site of the mutation (Y144) shows the presence of a surface-exposed negative charge that would be lost with the mutation. This would consequently alter prospective binding sites for neutralizing antibodies raised against the non-variant SARS-CoV-2 strains (Figure 3). The signature mutations that result in altered molecule surface character should therefore be noted for their potential effect on relevant partner protein interactions. H1101Y represents a similar type of mutation, where the surface character of the Spike protein is altered. The replacement with Tyrosine at the 1101 position provides a positive charge on the surface in 3 of the 5 possible rotamers conformations. The remaining two conformations retain an interaction with an existing contact for H1101 (i.e.F1103), with the addition of another contact (N1098) (Figure 4). The different conformations available for Y1101 therefore provide two possible implications for the mutation. First, for altered surface character, and second for increased stability for the Spike protein monomer. Both implications may be utilized for the advantageous evolution of the virus, rationalizing the observation of these types of mutations in the emergent Philippine variant.

**Figure 3.**
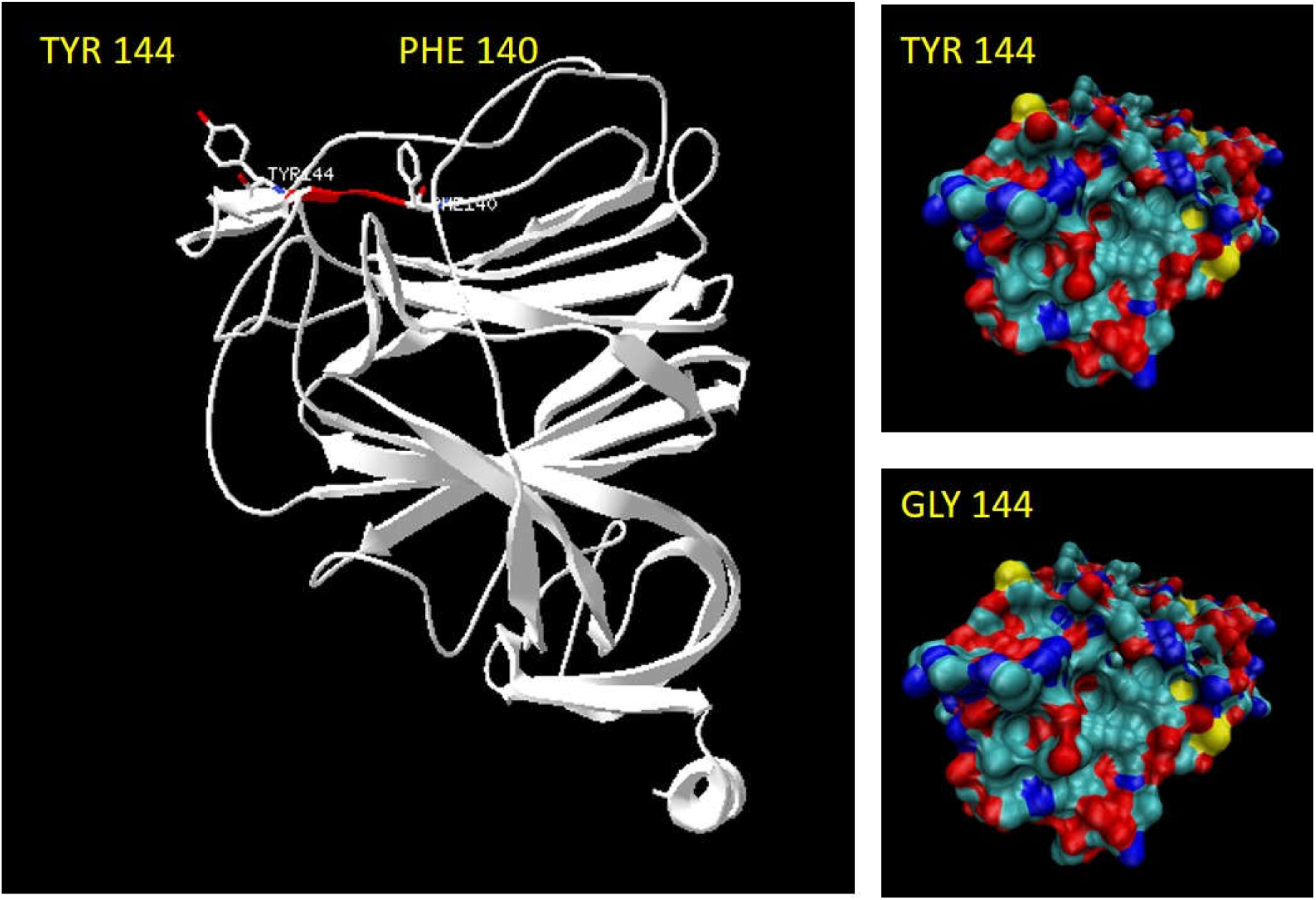
LGV141-143del implications on NTD structure. The removal of the LGV tripeptide sequence from the N-terminal domain may alter structural stability and surface characteristics. The LGV sequence, colored red in the ribbon diagram for the NTD structure, is flanked by aromatic residues (F140 and Y144). A deletion of the LGV tripeptide will shift the position of these aromatic residues and may alter the stability and surface characteristics of the NTD. Movement of the P140 residue may provide additional intramolecular contacts between itself and other aromatic residues thereby stabilizing NTD structure. A shift in the Y144 position can also alter the surface charge properties. A Y144del mutation previously reported for the UK variant (B.1.1.7) is approximated here by a G144 model. The loss of the negative Y144 sidechain alters the electrostatic map of the NTD surface, potentially inhibiting its recognition by neutralizing antibodies.

**Figure 4.**
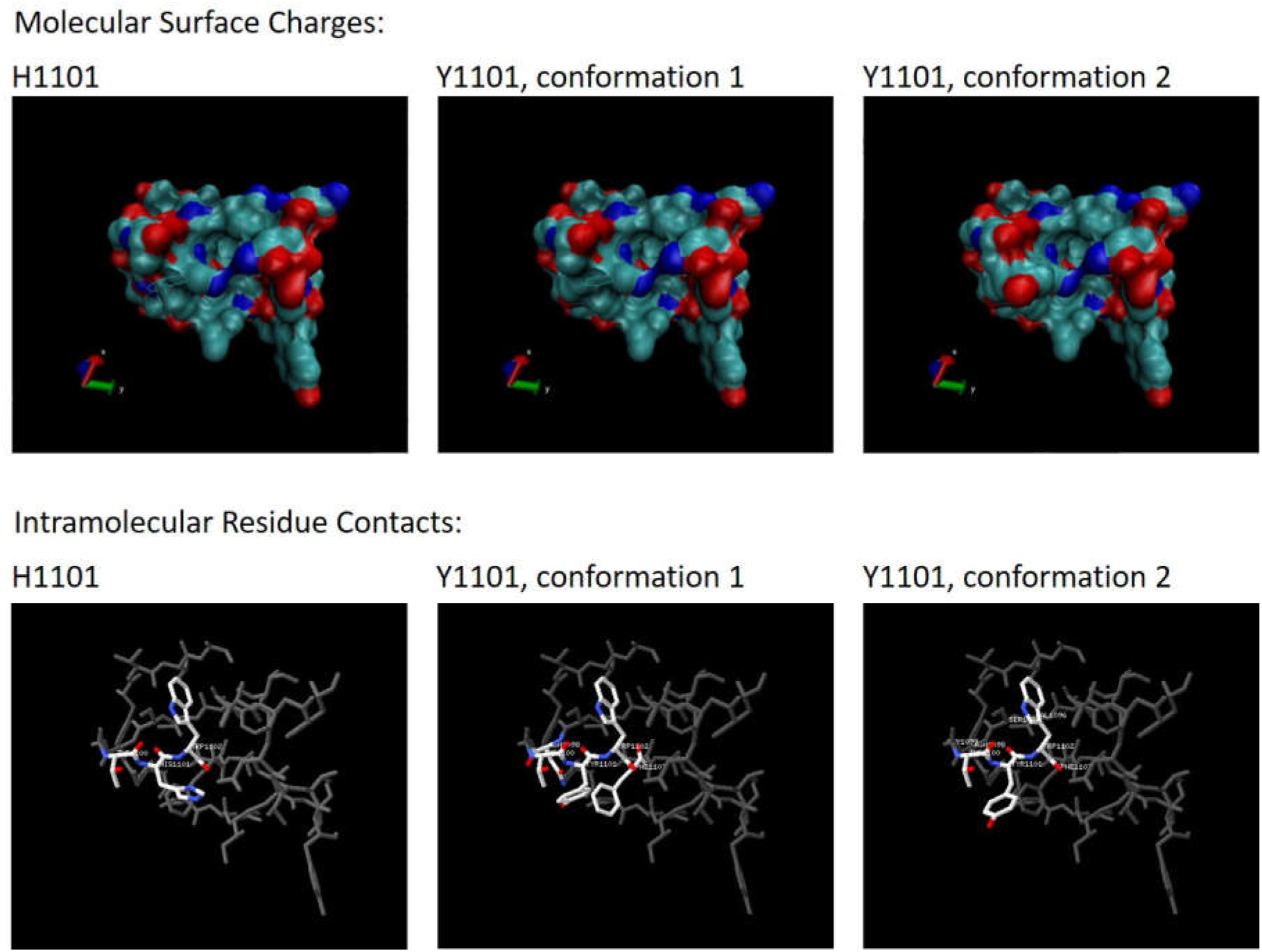
Structural Implications of the H1101Y mutation The H1101Y mutation is predicted to alter the RBD-FP linker domain of the Spike Monomer in two possible ways. In 2 of the 5 possible rotamer positions of the substituted Tyrosine (Y1101, conformation 1), increased contacts are formed with the neighboring residues (≤ 3Å). Most notably a pi bond with F1103 is added which would stabilize this structure. In the other 3 rotamer positions (Y1101, conformation 2), Y1101 adopts an exposed conformation that alters the electrostatic surface of the molecule, with the addition of a negative charge.

### Significance of the three amino acid L141-V143 deletion on Spike protein monomer stability

Similar to H1101Y, several of the other observed mutations may be associated with effects on the stability of the Spike-protein monomer. L141-V143 occurs on the surface of the N-terminal domain (NTD). These residues are flanked residues with aromatic side-chains (e.g. F,H,P,Y,W). Models were generated for the isolated NTD of the Spike Protein using RaptorX (Källberg et al. 2012) and results for the wildtype NTD sequence, and LGV141-143del variant were compared. The models for the wildtype sequence were observed to adopt a more extended structure for the L141-V143 location. Pi bonding between aromatic residues were restricted to the regions flanking the LGV141-143del location (Figure 5 B, C). Deletion of LGV141-143 brings these flanking aromatic residues into closer proximity, allowing a more compact structure for the NTD, stabilized by multiple pi-bonding interactions (Figure 5 E, F). Increased stability for the NTD, as with other structural domains of the Spike protein, may serve to prolong effective interaction with the ACE II receptor, and ultimately promote viral infection.

**Figure 5.**
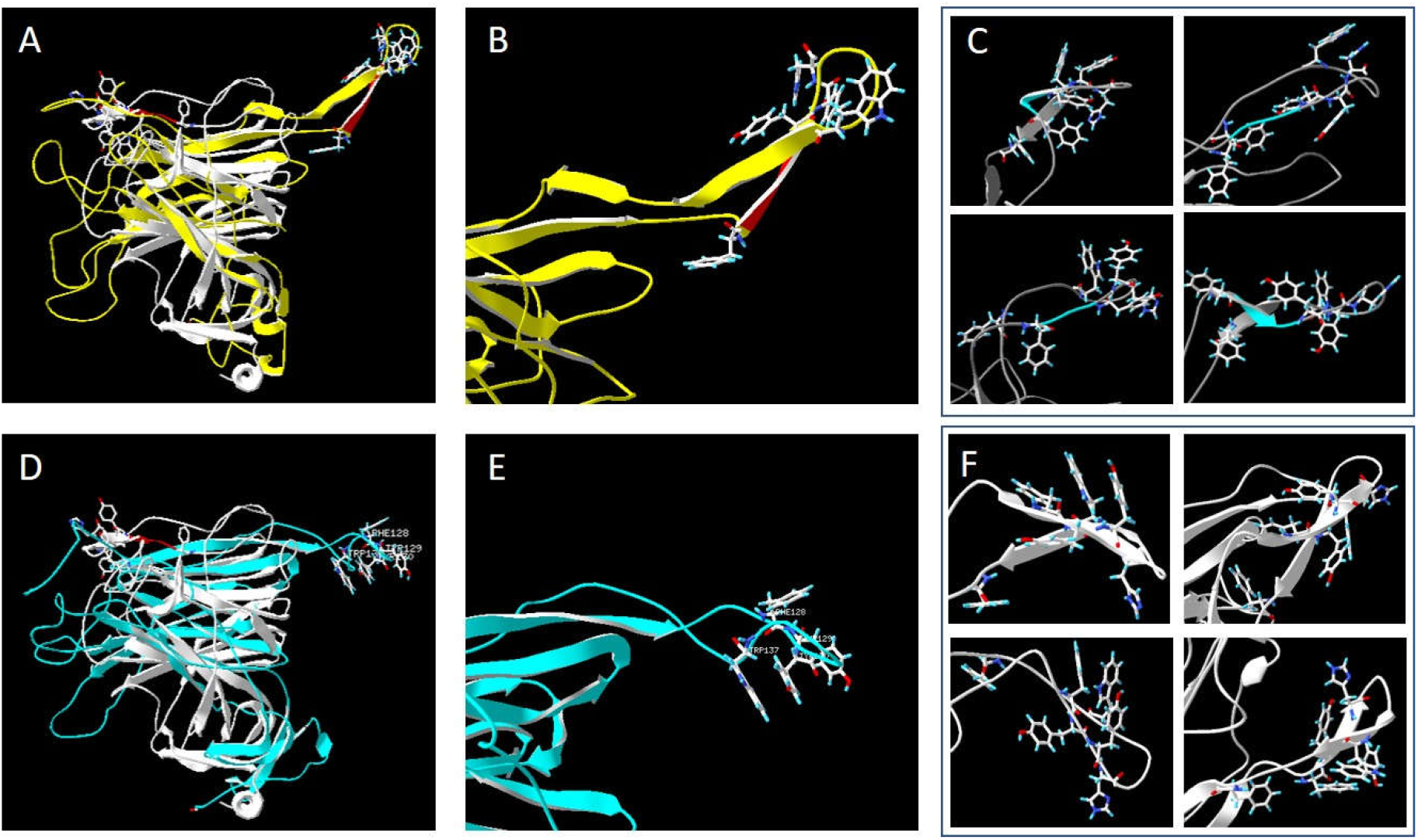
LGV141-143del implications on NTD structure. The removal of the LGV tripeptide sequence from the N-terminal domain may alter structural stability and surface characteristics. Models for the wildtype NTD, and the LGV141-143del mutant NTD were generated using RaptorX (Källberg et al, 2012). (A,D) The structures for these models (Yellow, and Cyan, respectively) were fit against the iTASSER generated NTD structure (S_ACE2.pdb, Zhang Lab 2020, white). The LGV sequence is highlighted (red in A,B; cyan in C) in the ribbon diagrams for the NTD. The models for the RaptorX generated models are observed to vary from that of iTASSER in terms of the position of the LGV containing loop. (B,D) However, focusing on the modelled structural effects of the LGV141-143del, the removal of the LGV sequence is observed to result in a more compact structure, stabilized by pi-bonded aromatic residues (Y130 – W137). (B, C) Aromatic residue interactions are were restricted to residues flanking the LGV sequence, resulting in an extended conformation for the structure in 4 of 5 returned models. (E,F) LGV141-143del results in a more condensed structure for the section, with stabilizing aromatic residue interactions in 5 of 5 returned models.

Altered monomer residue interactions are also observed for the P681H mutation in the RBD-FP linker. The modelled structure for the Spike protein trimer, with one monomer bound with the ACE II receptor (S_ACEII.pdb; iTASSER model; https://zhanglab.ccmb.med.umich.edu/COVID-19/) shows two (2) distinct conformations for the region containing P681 (Figure 6A). Within the same trimer, if the monomer is bound by an ACE II receptor, P681 exists within an extended coil (Figure 6B). In the absence of the bound ACE II receptor, the region proximal to P681 is stabilized into antiparallel beta strands (Figure 6C, D). The different conformation attained with ACE II receptor interaction suggests the involvement of allosteric effects that may be linked to viral function. Interestingly, P681 is directly adjacent to the documented furin binding sequence (RRAR) found in the Spike protein (Coutard et al. 2020). Stability of the FCS may be increased with additional interactions made between H681 and R683. These additional contacts were seen for 5 out of 8 possible rotamers for the introduced Histidine residue. Increased contact between H681 and R683 would stabilize the canonical FCS and allow more efficient recognition and cleavage.

**Figure 6.**
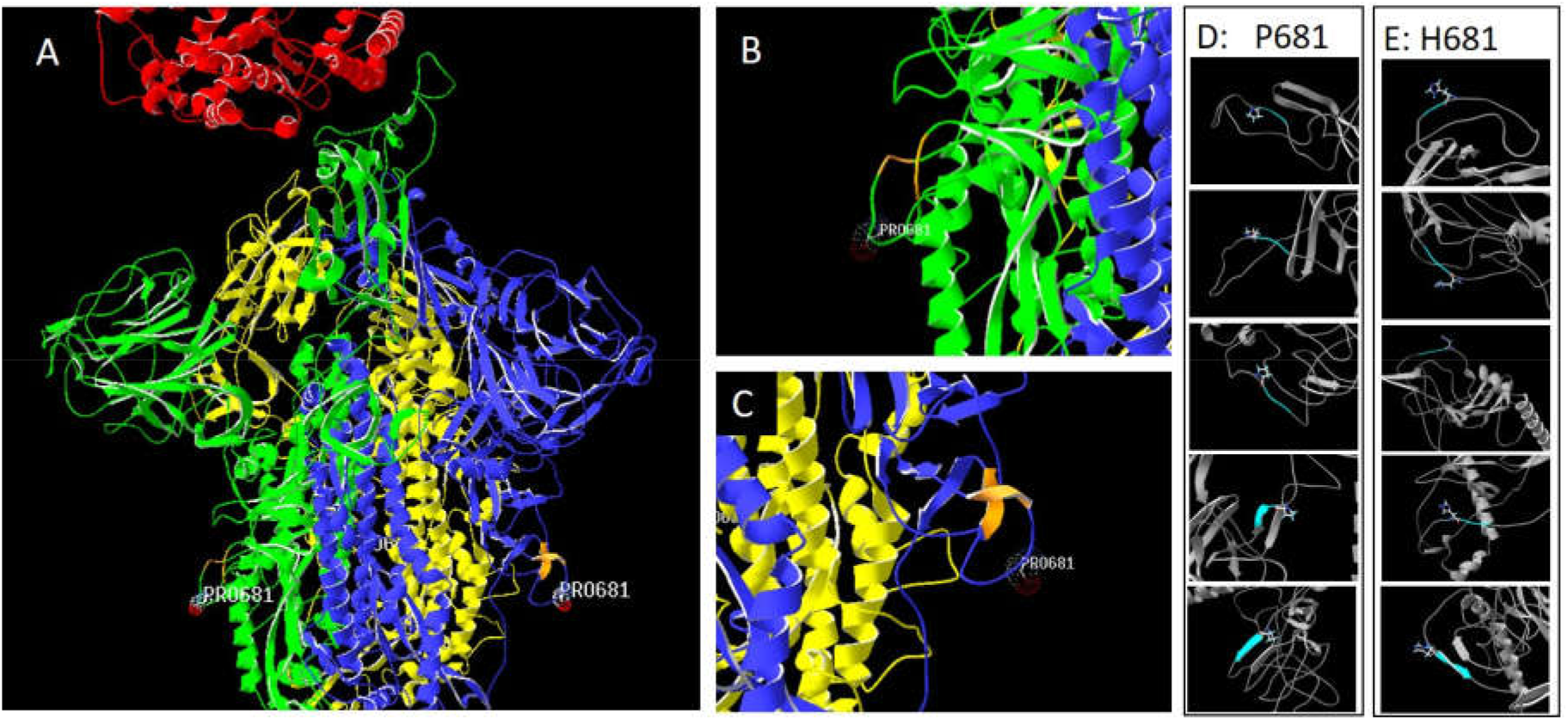
Structural Implications of the P681H mutation. The P681H mutation is located in the region between the Spike RBD and FP domains. (A) This linker region is observed to exist in two conformations in the iTASSER modelled spike protein bound to the ACE II receptor. (B) The ACE II receptor bound monomer (Chain C) exhibits an extended coil structure at this location. (C) Monomers without bound receptor (Chains B and A) show a more stabilized conformation, with proximal anti-parallel beta strands. (D) The stabilized conformation was observed in 2 out of 5 models for the wildtype protein sequence with P681. (E) Substitution with Histidine (i.e. H681) results in the stable conformation in only one out of the 5 modelled structures.

The existence of the furin cleavage site (FCS) has been shown to increase SARS-CoV-2 infectivity compared with SARS (Xia, et al, 2020). The extended coil conformation adopted near P681 with ACE II receptor binding suggests a receptor-induced conformational change in the FCS that would also potentially favor cleavage and facilitate more efficient viral fusion with the membrane of the target cells.

Replacement of P681 with Histidine was shown to result in more extended coil conformations for the P681H variant. Relating this to the observed effects with ACE II receptor binding, the P681H variant seems to stabilize the extended conformation assumed with the receptor-bound state (Figure 6E), possibly achieving this conformation even in the absence of receptor binding. This could allow for more efficient furin cleavage of the Spike protein and thus promote viral entry into the cell. P681H would also result in a slight increase in the polar character of the surface, potentially increasing stability in aqueous environments, and altering neutralizing antibody binding properties.

The P681H mutation has also been identified in several other variants, including the one from the UK. A recent study which is currently under review posits that this particular mutation is emerging worldwide (Maison et al., preprint). The actual biological effects of the mutation, however, remain unknown. It is commonly believed that alterations in the sequences within and surrounding the furin cleavage site are often associated with coronavirus evolution, particularly when moving from one host to another (Gallagher and Garry, 2020), and the apparent convergent evolution occurring at this residue in multiple, unrelated samples indicates that this mutation is likely to confer an evolutionary advantage to the virus. A recent study by Johnson, et al. (2021) showed that the deletion of P681 along with R682, R683, and A684 resulted in the reduced cleavage of the viral Spike, altered viral infectivity in cell culture, and reduced lung pathogenicity in a hamster model of SARS-CoV-2 infection. It also led to decreased neutralization by immune sera from patients infected with WT virus. This reinforces the hypothesis that alterations of the Spike protein at and around P681 may affect viral function and fitness.

In addition to increased residue contacts that stabilize the Spike protein monomers, some of the observed signature variations resulted in additional contacts between residues from different monomers, and therefore stabilize the trimeric form. Increased inter-monomer contacts are seen with the E1092K mutation, where all three (3) monomers are linked. The introduction of the longer Lysine sidechain allows additional interactions between K1092 of 1 monomer (i.e., Chain C) with W886, and Y904 of the next monomer (i.e., Chain B), and also with F1101 of the third (i.e. Chain A) (Figure 7A, B).

**Figure 7.**
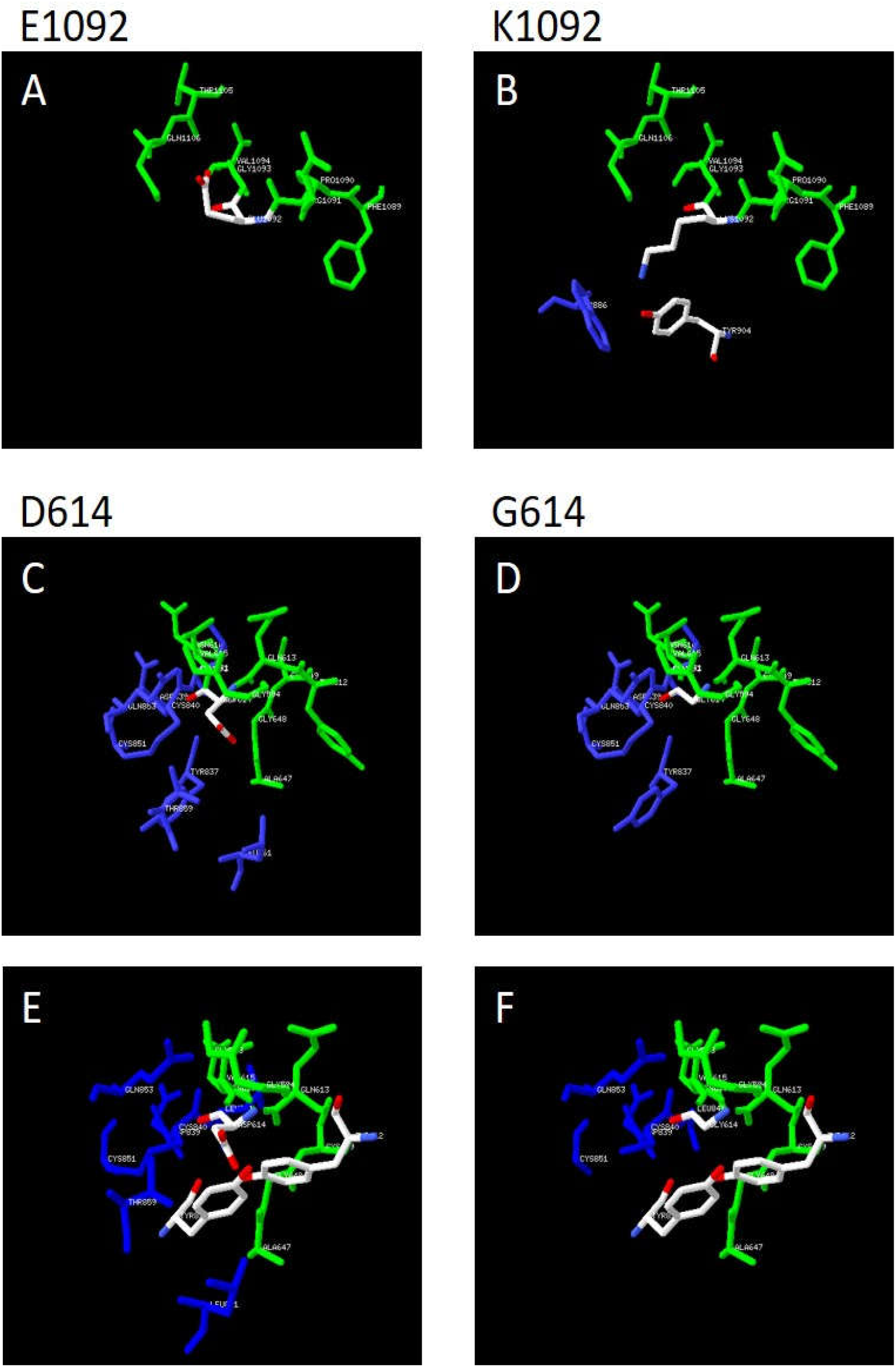
Changes in Spike protein monomer linkages The D614G and E1092K were observed to alter interactions between neighboring Spike protein monomers. Residues in chain C are colored green, and residues in chain B are colored blue. (A,B) Potential interactions within ≤ 5Å for E1092 were only found within the same monomer (Chain C). The substitution of K1092 increased the potential interactions to include residues W886 and Y904 in the next monomer (Chain B). Interestingly, K1092 provides a complimentary charge to Y904, and Y904 may interact with W886 through their aromatic rings, likely stabilizing the monomer structure at this domain. (C,D) D614’s location in the Spike protein allows it to have interactions with residues from its own monomer and the next (Chain C, and B). The substitution with G614 leads to a decrease in contacts with the next monomer. (E,F) The shorter side chain of G614 may influence the contacts between the Spike protein monomers. Rotation of Y687 from one monomer (Chain B), and Y612 from another (Chain C) may allow the formation of an inter-monomer contact. While this contact may occur with both D614 and G614, the similar charges for D614 and Y612 may cause repulsive charges that would inhibit its formation.

The relevance of these increased inter-monomer contacts may be associated with the documented increased transmissibility conferred by the D614G mutation (Hou et al. 2020; Yurkovetskiy, et al. 2020; Zhang L et al. 2020) This mutation is located in the RBD-FP connecting two monomer structures (Chain C and B in the S_ACEII.pdb model, https://zhanglab.ccmb.med.umich.edu/COVID-19/). Similar to LGV141-143del, removal of intervening structures, provide additional contacts between proximal residues with charge/character compatibility. In this case, replacing the negatively charged sidechain of D614 with a Hydrogen (G614), results in less stearic hindrance for the association of Y612 and G648 from one monomer, with Y837 of another (Figure 7F).

The involvement of aromatic residue interactions in stabilizing the trimeric structure may be observed in their increased presence for areas wherein the individual monomer chains condense into more compact structures. Figure 8 shows the location of all aromatic ring containing residues (W,Y,F,H,P) within the Spike protein trimer. These are found to exist only in the condensed areas of the structure, consistent with the hydrophobic effect. The only condensed area without these residues is near the assigned transmembrane region from AA 1214-1237 (Huang et al. 2020). W1214, Y1215, W1217 do exist in this region at its N-terminal end. Given these findings, it is likely that a V1175F mutation, placing aromatic residues in the HR2 domains of the monomers, would provide an additional anchor point between Spike protein monomers and further stabilize the trimeric structure.

**Figure 8.**
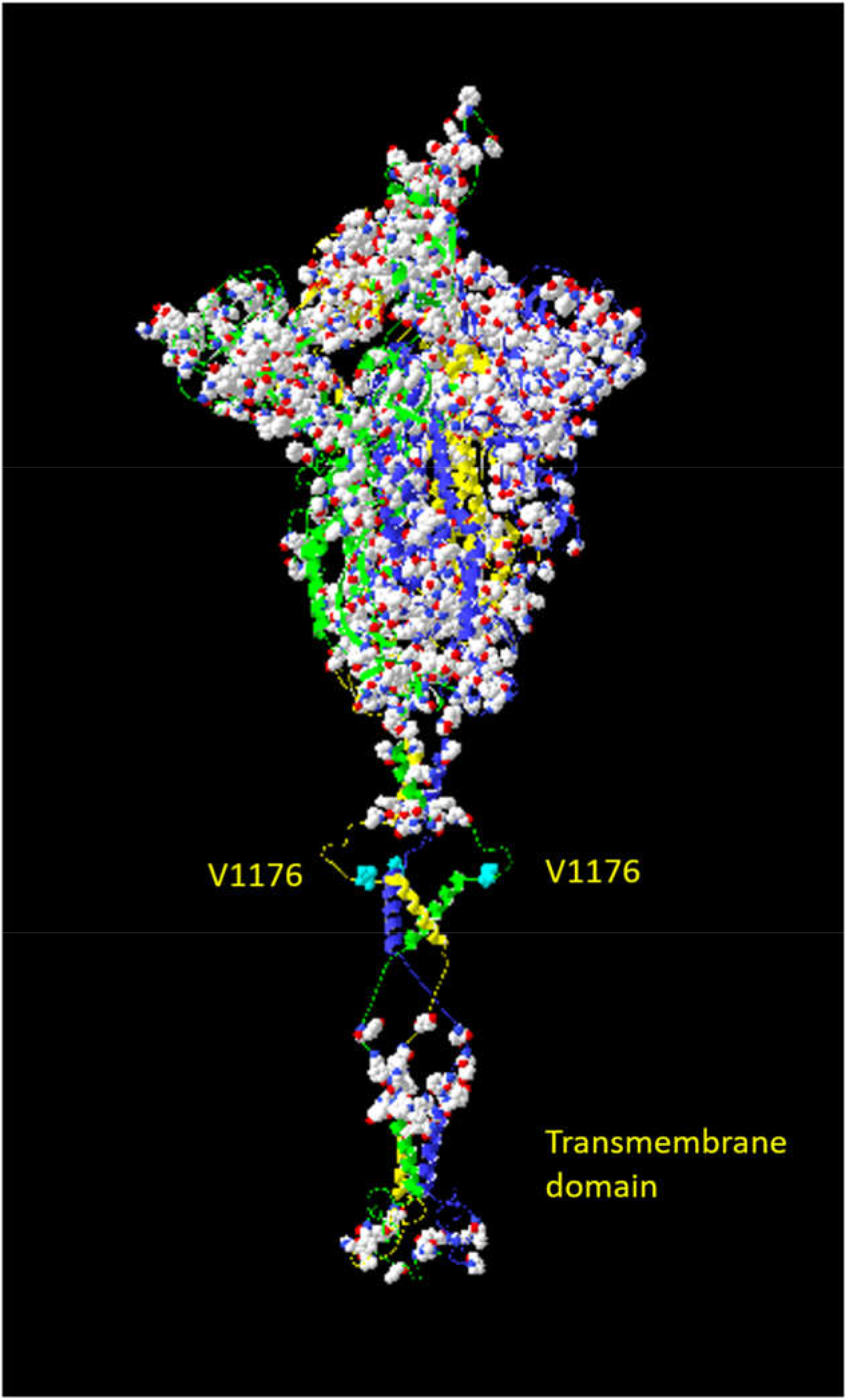
Aromatic residue positions within the Spike protein trimer. All residues with aromatic sidechains (i.e. W,Y,F,H,P) were represented as space-filling models. Their location in all condensed regions of the Spike protein structure (except the transmembrane domain) coincides with their observed involvement in stabilizing these domains through pi bonding. V1176 is colored cyan and is found outside these condensed areas. A V1176F substitution will introduce an aromatic residue to this region and allow an additional stabilizing interaction for the Spike protein monomers.

## CONCLUSION

### Insights for epidemiological investigation

In this study, we analyzed *in silico* the potential structural effects of multiple mutations on the Spike protein detected in viruses belonging to the P.3 lineage from Philippine samples. These samples were collected mainly from the Central Visayas region between January 30, 2021 and February 2, 2021, in the context of a reported increase in case numbers in this area in the first two (2) months of 2021 (COVID-19 Tracker | Department of Health website (doh.gov.ph)).

The predicted structural effects of the documented mutations rationalize their significance for the evolution of a potentially more transmissible variant of SARS-CoV-2. The recent rise in cases in the region from which these signature mutations were identified supports this hypothesis. The emergence of variants that are better able to infect, or bypass host defenses is a natural part of viral evolution. The key therefore is to (1) mitigate the potential effects of the increased transmission of the virus through continued observation of the recommended minimum public health practices such as social distancing, wearing of masks and face shields and other protective equipment, frequent handwashing, avoidance of crowds, etc., and to (2) identify the mechanisms through which the SARS-CoV-2 virus has, so far, successfully improved its capacity for infection and survival, and then use this information to determine ways to inhibit the virus and to help anticipate and preempt the effects of any future mutations that may naturally occur during the course of its evolution.

This study has shown that SARS-CoV-2 evolution has utilized modifications that increase receptor binding, and molecular interactions within the monomer and trimer structures. In the P.3 variant, these changes are believed to involve increased contacts between aromatic-ring containing residues and potential salt-bridge formation (E-K contacts). Increased contacts are achieved in several ways. Some involve the addition of a new aromatic residue (i.e. N501Y, P681H, H1101Y, V1176F). Others involve the removal of intervening structures, allowing new contacts to be made (i.e. LGV141-143del; D614G). In addition, the introduction of Lysines, with their long side chains, was also observed to allow increased residue contacts, usually through salt-bridge formation. The E484K mutation resulted in additional long-range contacts between the Spike protein and the ACE II receptor, and the E1092K mutation allows for increased contact for all three chains of the Spike protein trimer. It must be noted that the observed mutations were distributed across the Spike protein structure, and are likely to result in compounded effects for increasing the Spike protein’s stability, and altering its receptor interactions.

Although many of the mutations reported herein have been documented before, this is, to our knowledge, the first report of the N501Y, E484K, P681H and N-terminal LGV141-143del mutations occurring together in a single virus. Figure 9 illustrates the signature mutation positions within the emergent Philippine variant with the expected functional effects of these mutations, and how these relate to the locations of mutations in other emergent variants worldwide. The co-existence of these mutations of potential clinical significance, along with several others within the P.3 lineage suggest the evolution of the virus into a possibly more transmissible, and less neutralizable form. This emphasizes the need for its further characterization in terms of transmission dynamics, pathogenicity and susceptibility to existing vaccines and antibody treatments.

**Figure 9.**
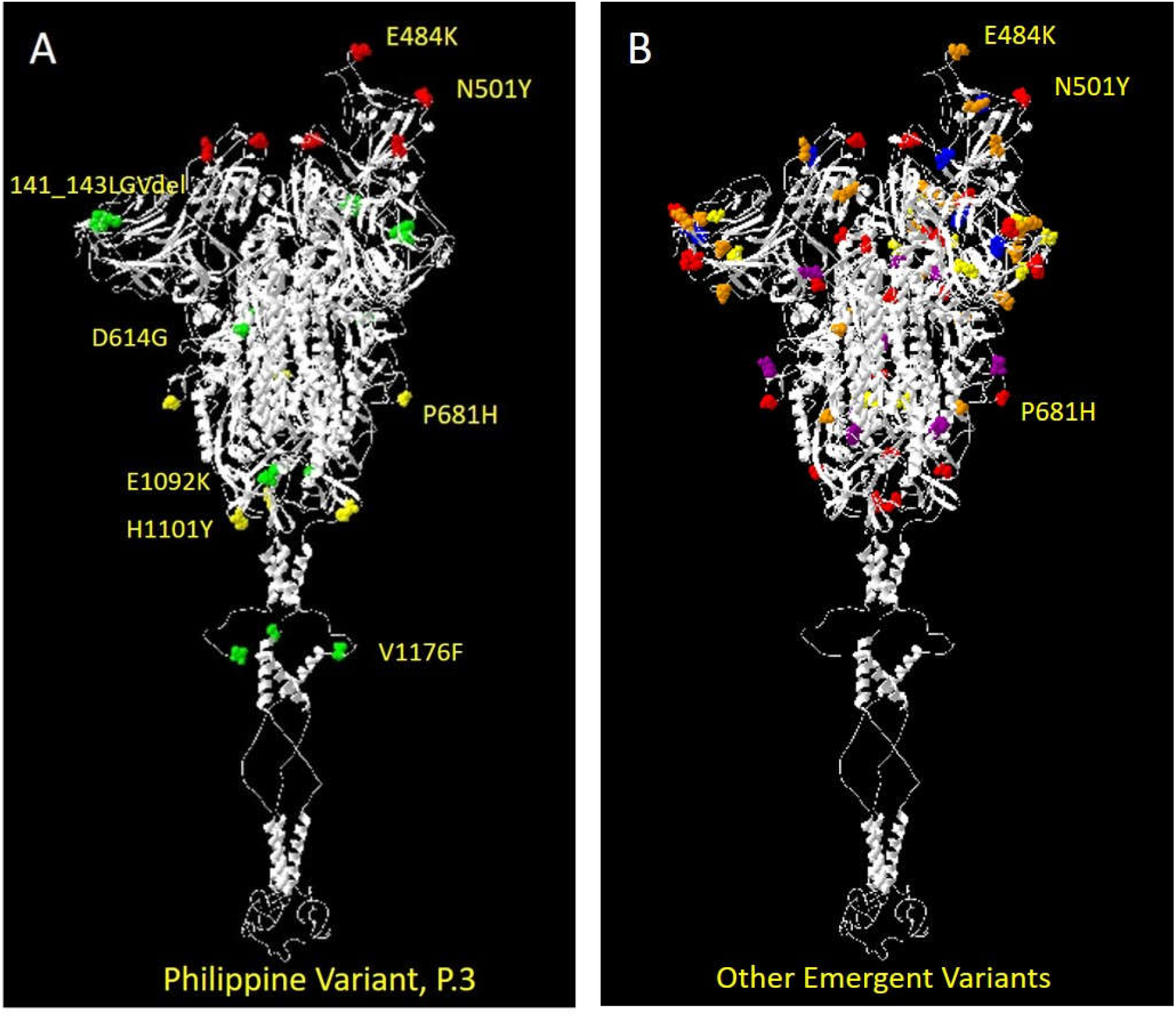
Location of Mutations in Several SARS-CoV-2 Variants (A) Position of Signature Mutations for the Philippine Variant (P.3). Mutations are colored based on their predicted functional effect:

a. Red: Affects ACE II receptor interactions;
b. Yellow: Affects molecular surface character;
c. Green: Increased monomer/trimer interactions. (B) Position of Signature Mutations in several reported SARS-CoV-2 Variants:

a. Red: B.1.1.7 (UK Variant)
b. Orange: B.1.351 (South African Variant)
c. Yellow: P.1 (Brazil Variant)
d. Blue: B.1.429 (CA Variant)
e. Violet: B.1.526 (NY Variant)

## ACKNOWLEDGMENTS

The authors acknowledge the contributions of the various COVID-19 testing laboratories in Central Visayas and the other members of the Genomic Biosurveillance Network of the Department of Health, Philippines, for sharing their samples; the members of the DNA Sequencing and Bioinformatics Core Facilities of the Philippine Genome Center; and the Epidemiology Bureau of the Department of Health, for making the P.3 sequences publicly available.

## NOTES ON APPENDICES

None

